# Antibody escape and cryptic cross-domain stabilization in the SARS-CoV-2 Omicron spike protein

**DOI:** 10.1101/2022.04.18.488614

**Authors:** Kamyab Javanmardi, Thomas H. Segall-Shapiro, Chia-Wei Chou, Daniel R. Boutz, Randall J. Olsen, Xuping Xie, Hongjie Xia, Pei-Yong Shi, Charlie D. Johnson, Ankur Annapareddy, Scott Weaver, James M. Musser, Andrew D. Ellington, Ilya J. Finkelstein, Jimmy D. Gollihar

## Abstract

The worldwide spread of severe acute respiratory syndrome coronavirus 2 (SARS-CoV-2) has led to the repeated emergence of variants of concern. The Omicron variant has two dominant sub-lineages, BA.1 and BA.2, each with unprecedented numbers of nonsynonymous and indel spike protein mutations: 33 and 29, respectively. Some of these mutations individually increase transmissibility and enhance immune evasion, but their interactions within the Omicron mutational background is unknown. We characterize the molecular effects of all Omicron spike mutations on expression, human ACE2 receptor affinity, and neutralizing antibody recognition. We show that key mutations enable escape from neutralizing antibodies at a variety of epitopes. Stabilizing mutations in the N-terminal and S2 domains of the spike protein compensate for destabilizing mutations in the receptor binding domain, thereby enabling the record number of mutations in Omicron sub-lineages. Taken together, our results provide a comprehensive account of the mutational effects in the Omicron spike protein and illuminate previously unknown mechanisms of how the N-terminal domain can compensate for destabilizing mutations within the more evolutionarily constrained RBD.

## Introduction

The continuous evolution and spread of severe acute respiratory syndrome coronavirus 2 (SARS-CoV-2) has produced variants of concern (VOCs) and variants of interest (VOIs) with enhanced immune evasion, transmissibility, and occasionally increased disease severity^1–5^. Omicron (or B.1.1.529 in the PANGO nomenclature) rapidly displaced the Delta VOC globally^6–8^. The BA.1 sub-lineage of Omicron caused record numbers of infections and breakthrough cases in fully vaccinated and previously infected individuals^9,10^. As of February 2022, the BA.2 lineage has displaced BA.1 in many countries and shows additional enhanced transmissibility over all prior VOCs^11,12^.

The SARS-CoV-2 spike protein is key to both transmissibility and immune evasion^13^. This homotrimeric protein is displayed on the SARS-CoV-2 capsid surface, mediates virus binding and entry into host cells, and elicits a strong immune response that gives rise to neutralizing antibodies and a robust T-cell response^14–16^. The spike protein ectodomain (ECD) is the primary immune target and consists of three main functional units: the N-terminal domain (NTD), receptor binding domain (RBD), and the fusogenic stalk (S2)^17^. Because of its importance in cell entrance and immune escape, spike mutations accumulate rapidly in circulating viral variants (**Figure 1A**). The NTD appears to tolerate the most mutations, harboring 31% of all amino acid substitutions and 84% of indels found in circulating variants (GISAID database accessed on 18/December/2021) (**Figure 1B**), while the RBD and S2 regions are more restricted in the structural changes that they can tolerate, likely due to conserved functional constraints of host-cell receptor binding and membrane fusion. The functional consequences of most of these mutations— and the molecular epistasis of multiple mutations—is key for understanding viral evolution and interactions with our immune system.

**Figure 1:**
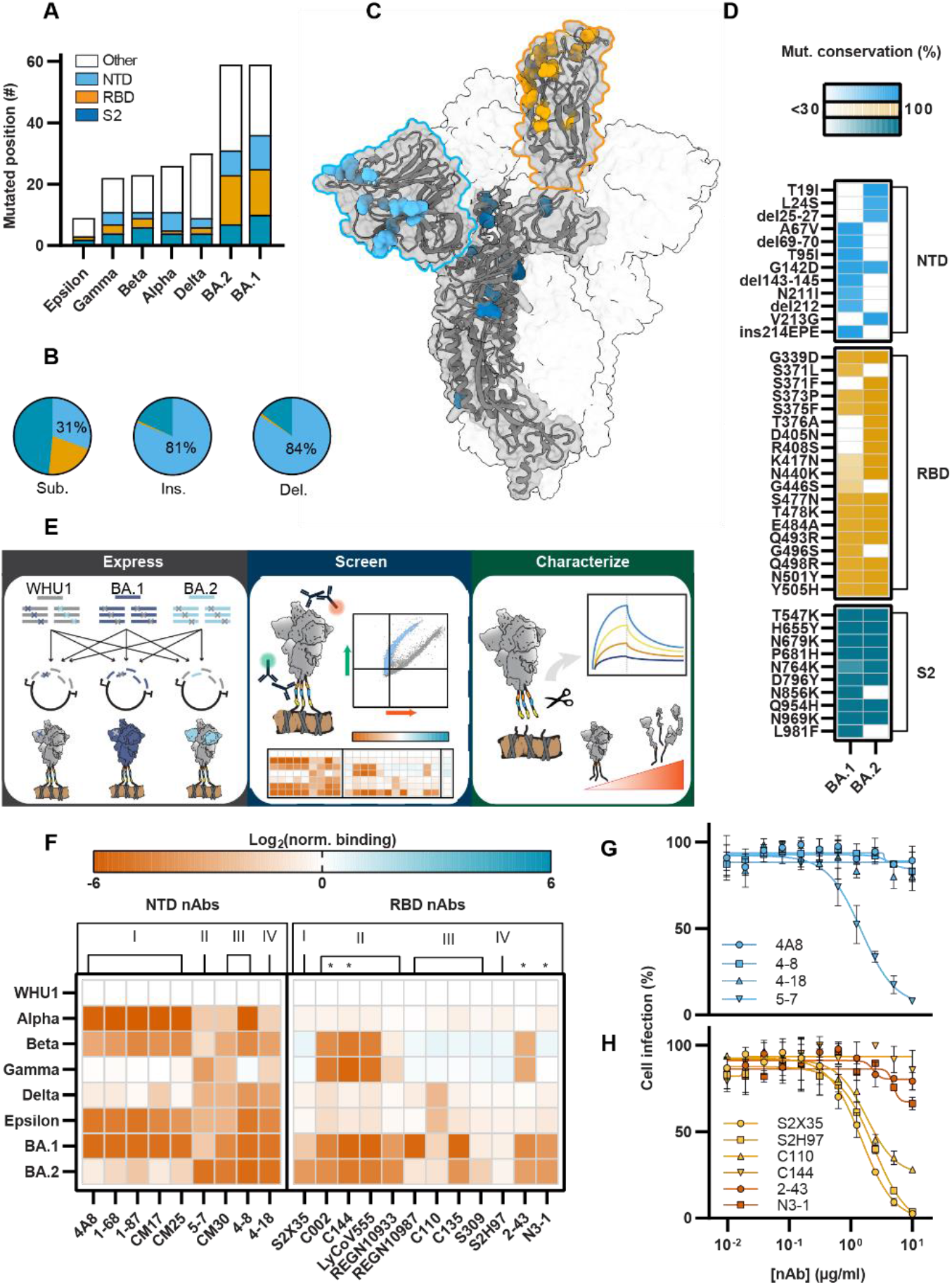
Omicron spike protein ectodomain has dozens of mutations contributing antibody escape. A. The number of mutated amino acid positions for each VOI and VOC. Spike NTD mutations, light blue; spike RBD mutations, yellow; spike S2 mutations, dark blue; other mutations, white. B. Distribution of all non-synonymous mutations (substitutions = 42,077,816; insertions = 31,063; deletions = 15,664,146, colored as in Figure 1A) found in GISAID (accessed on 18/December/2021). The NTD has the majority of insertions and deletions (81% and 84% respectively). C. SARS-CoV-2 spike ectodomain structure (PDB: 7DDN^59^) with mutations found in BA.1 and BA.2 colored by domain as in Figure 1A. D. Mutations found in the Omicron spike variants. Shading indicates the percentage of BA.1 or BA.2 strains containing these mutations, as analyzed on outbreak.info (accessed on 12/February/2022). E. Spike Display overview. Spike protein ectodomains with different mutation sets are constructed using a semi-automated Golden Gate-cloning pipeline, displayed on the surface of HEK293T cells, and assayed in high-throughput with flow cytometry. Biophysical characterization is performed with spikes cleaved from cell surfaces. F. Relative monoclonal antibody binding to spikes from VOCs (Alpha, Beta, Gamma, Delta, and Omicron) and a VOI (Epsilon). Red, decreased binding; blue, increased binding; normalized to the original WHU1 spike with D614G mutation (top row). Mean ± SD of log-transformed values from at least two biological replicates. Spike domain targets and epitope classifications of antibodies shown on top, * = quaternary binding. G. Authentic BA.1 virus neutralization for selected NTD-directed monoclonal antibodies. Mean ± SD of two biological replicates. Curves are a Sigmoidal (4PL, X), Least squares fit, IC50 values listed in Figure S1E. H. Authentic BA.1 virus neutralization for selected RBD-directed monoclonal antibodies as in Figure 1G.

Omicron BA.1 and BA.2 sub-lineages have an unprecedented 33 and 29 nonsynonymous changes relative to the ancestral Wuhan-Hu-1 (WHU1) lineage. These include four distinct amino acid (AA) deletions (del), one insertion (ins), and 36 substitutions distributed across the ECD (**Figure 1C**). BA.2 shares 21 of the mutations in BA.1, but also contains eight unique substitutions and deletions (**Figure 1D**). Some of these mutations increase the evasion of neutralizing monoclonal antibodies (mAb), rendering most mAb therapies ineffective^18^. Similarly, the mutations significantly reduced authentic virus neutralization by convalescent and vaccinated/boosted sera^19,20^. While recent studies have probed small numbers of individual amino acid changes within BA.1 and BA.2^21,22^, questions about the synergistic and contextual effects of most Omicron mutations remain unanswered.

Here, we leverage mammalian cell surface display of the spike protein to rapidly characterize the expression, antibody binding, and cell receptor affinity of coronavirus spike ECDs^23^. We characterize the effects of all Omicron mutations with respect to human ACE2 (hACE2) binding, spike protein stability, and escape from multiple distinct classes of mAbs. We compare the effects of individual spike protein mutations between the WHU1 and Omicron mutational contexts to reveal how these mutations alter antigenicity and hACE2 affinity. These results also explain how Omicron evades neutralizing mAbs, including those with quaternary binding between adjacent spike protomers, via changing both the surface epitopes and RBD conformational dynamics. Finally, we show that NTD mutations potentiate new RBD mutations, expanding the ability of the RBD to further evolve under increased evolutionary pressure from our adaptive immune response.

## Results

### Omicron spike proteins have distinct antigenic features

We used mammalian cell surface display to compare the antigenicity and expression of the BA.1 and BA.2 ECDs (residues 1-1208) to earlier VOCs (**Figure 1E**). We transiently express spike variants on the surface of human embryonic kidney (HEK293T) cells. Surface-displayed spikes are then immunostained to measure expression, antibody binding, and hACE2 affinity by two-color flow cytometry (**Figures S1A-B, see Methods**)^23^.

We first cloned spike proteins from five VOCs (Alpha, Beta, Gamma, Delta, and Omicron BA.1 & BA.2) and one VOI (Epsilon), representing the dominant variants during different surges of the pandemic. The globally-dominant D614G mutation and six prefusion stabilizing prolines were incorporated into all variants to increase surface expression and maintain the prefusion conformation^24–26^. We assayed each variant for expression and antibody escape potential with a set of 21 mAbs, many with known high-resolution structures (**Figures 1F and S1C-D**)^27–29^. Of these, nine are NTD-targeting mAbs^28,30,31^, which we previously classified based on their binding epitopes (classes I-IV) ^23^. The remaining 12 target all four classes of neutralizing RBD epitopes^27,32^. We include the clinically-used REGN10933 (casirivimab) and REGN10987 (imdevimab)^33,34^, LY-CoV555 (bamlanivimab)^35^, and S309 (sotrovimab)^36,37^ antibodies. We also tested four mAbs with quaternary RBD binding modes^27,31,38^ (C002, C144, 2-43, and N3-1) and the pan-variant mAb S2H97^31,32,37,38^. Together, this panel provides a comprehensive overview of neutralizing antibody escape by variant spike proteins.

Consistent with previous reports^20–22,39,40^, BA.1 and BA.2 show enhanced antibody escape compared to all other variants (**Figure 1F**). BA.1 escapes the majority of antibodies in our panel, including nearly all classes of NTD binding mAbs tested here (the only exception is the class II antibody 5-7). BA.1 spike is also refractory to many RBD targeting mAbs, with strong escape from class II binders, half of the class III binders, and all quaternary binders. BA.2 shows considerable antibody escape but remains susceptible to the class I NTD binders and some class III RBD antibodies. In contrast, BA.2 escapes 5-7, the noncanonical class I RBD binder S2X35, and the class III RBD binder S309 to a greater degree than BA.1.

To test whether these results translated to live virus, we performed microneutralization assays with authentic BA.1 virus and a subset of ten mAbs with known neutralization of WHU1. We tested four NTD-binding mAbs, one from each of the four binding classes (**Figure 1G**). Consistent with the cell surface display results, BA.1 completely escaped neutralization by 4A8 (class I), 4-8 (class III), and 4-18 (class IV). Only the class II mAb 5-7 had a measurable neutralizing effect. Similarly, we tested a representative of each of the four classes of RBD-targeting mAbs and two additional quaternary binding mAbs (**Figure 1H**). Again, BA.1 escaped the same antibodies as in the mammalian cell surface display assay. In the aggregate, the data showed that our screening methodology recapitulated other *in vitro* and *in vivo* observations and that the BA.1 and BA.2 spike proteins are antigenically distinct.

### BA.1 evades NTD-targeting mAbs better than BA.2

We sought to investigate the molecular mechanism of mAb binding escape by Omicron, starting with the effects of the NTD mutations. All VOCs contain at least one mutation in the NTD that increases viral escape from NTD-targeting mAbs (**Figure 1F**)^2,41–43^. Compared to previous VOCs, BA.1 and BA.2 have the most mutated NTDs: 11 and 6 NTD mutations, respectively. The BA.1 NTD has four AA substitutions, three deletions (del69-70, del143-145, del211), and a novel insertion (ins214EPE) (**Figure 2A**). The BA.2 NTD has four AA substitutions and one deletion (del25-27). Several of these mutations are located in the intrinsically disordered NTD loops (N-loops) that comprise an antigenic supersite (**Figures S2A-E**) ^23,29,44^. Strikingly, BA.1 and BA.2 only share one G142D substitution, which also appears in the Kappa and Delta variants.

**Figure 2:**
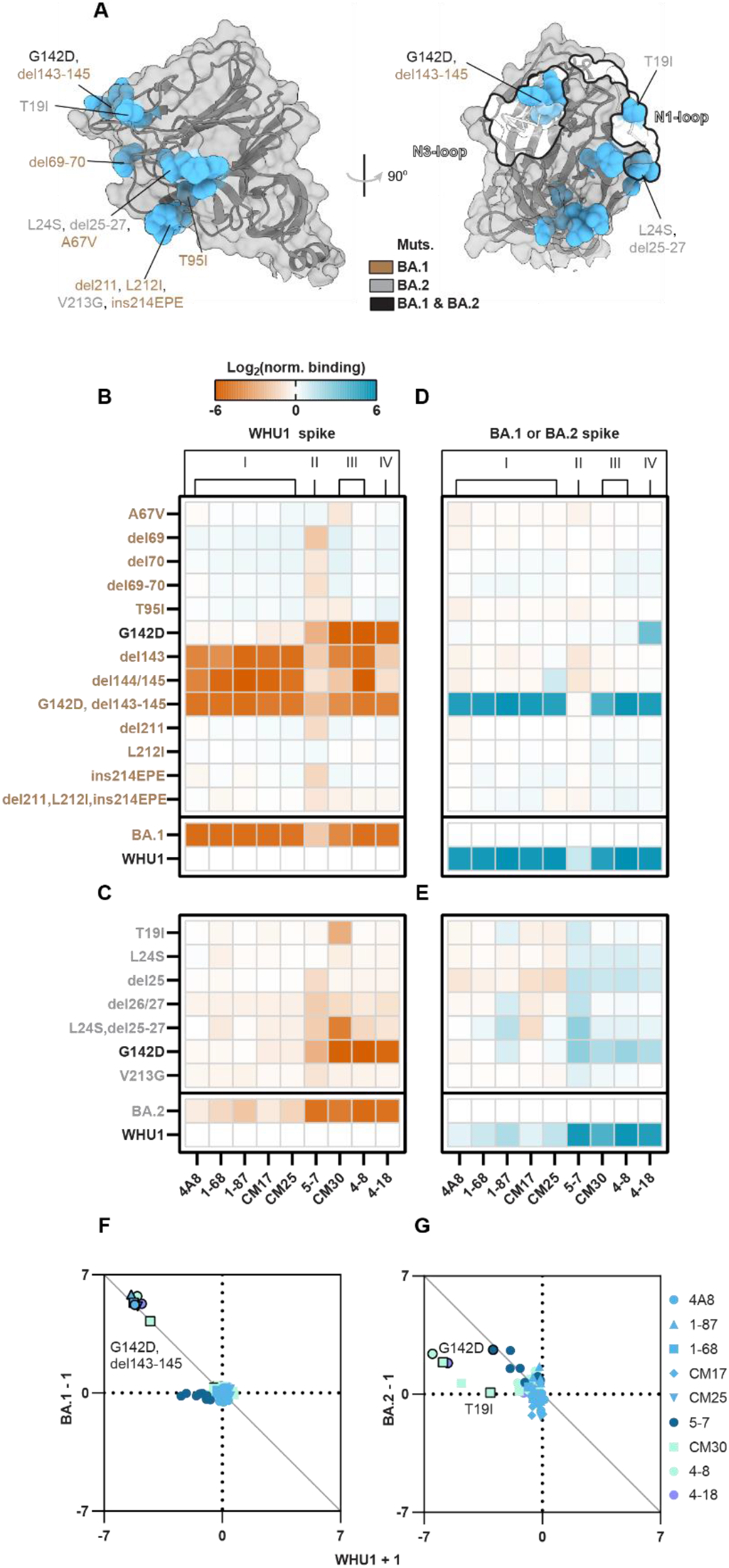
NTD indels and substitutions enable mAb binding escape. A. An enlarged NTD structure (PDB: 7DDN^59^) with nonsynonymous mutations from BA.1 (brown), BA.2 (grey), or both (black) indicated. B. Relative mAb binding to WHU1 spike proteins containing BA.1-NTD mutations. Red, decreased binding; blue, increased binding; normalized to the WHU1 spike. Mutations color coded as in Figure 2A. C. Relative mAb antibody binding to WHU1 spike proteins containing BA.2-NTD mutations. Colored as in B, normalized to the WHU1 spike. Mutations color coded as in Figure 2A. D. Relative mAb binding to BA.1 spike proteins containing reversions of the BA.1-NTD mutations to the WHU1 sequence. Colored as in Figure 2B, normalized to the BA.1 spike. E. Relative mAb binding to BA.2 spike proteins containing reversions of the BA.2-NTD mutations to the WHU1 sequence. Colored as in Figure 2B, normalized to the BA.2 spike protein. F. Comparison of log2(normalized binding) measurements for adding (+1) BA.1-NTD mutations to the WHU1 spike versus reverting (−1) the corresponding mutations from the BA.1 spike protein. Mutations with equal antigenic effects in both spike contexts are expected to fall on the diagonal line y = -x. G. Comparison of the effect of adding and reverting BA.2-NTD mutations as in Figure 2F. For all plots, mean ± SD of log-transformed values from at least two biological replicates

We first screened WHU1 spike protein variants containing each of the individual mutations found in BA.1 and BA.2 (**Figure S2F**)^23^. BA.1 completely escaped all NTD class I, III, and IV mAbs due to a series of contiguous mutations (G142D, del143-145) in the N3-loop (**Figure 2B**). Binding by 5-7 (class II), which interacts with the periphery of the NTD supersite, was reduced ~5-fold by these mutations^31,45^. The other BA.1 mutations (A67V, del69-70, T95I, del211, L212I, and ins214EPE) also had moderate effects on binding of mAb 5-7, but the other mAbs were not impacted.

To further interrogate these antigenicity changes, we dissected the mutation clusters into different combinations of mutations (**Figure S2G**). Class I mAbs, which predominately bind towards the apex of the N3-loop, had decreased binding due to a shortening of the loop (deletions) and the Y145D mutation (**Figure S2G-H**). Conversely, binding of class III and IV mAbs was more sensitive to mutations located at the base of the N3-loop (G142D and del142) (**Figure S2I**). Mutations del211 and ins214EPE each decreased binding of mAb 5-7 1.9-fold and moderately decreased pseudovirus neutralization (**Figures 2B and S2G**)^21^. These results document the importance of both the G142D substitution and the 143-145 deletion for the ability of BA.1 to escape different classes of NTD-targeting mAbs.

BA.2 still retained sensitivity to some NTD-targeting mAbs (**Figure 1F**). Mutations in the N1-loop (T19I, L24R, del25-del27) moderately decreased binding for class II, III, and IV mAbs (**Figure 2C and Figure S2J**) and in combination with G142D, BA.2 completely escaped antibody recognition at this site. Notably, the V213G substitution, found distal from the N-loops and in the same location as the del211, L212I, ins214EPE mutation cluster in BA.1, was inconsequential for mAb binding. However, these mutations may influence virus fitness at the RNA or protein level in ways that cannot be accessed by our platform.

Next, we tested whether BA.1 and BA.2 mutations impacted antibody escape at the NTD. Starting with each of the BA.1 and BA.2 sequences, we systematically reverted each of the NTD mutations to WT and measured the difference in mAb binding when compared with the full variant mutation set (**Figures 2D-E**). Reversion of the N3-loop mutations (G142D, del143-145) in BA.1 restored binding for all the affected mAbs (**Figure 2F**). However, reversion of the del211, L212I, ins214EPE mutation cluster failed to restore binding by mAb 5-7 to WHU1 levels. Reversion of N1-loop mutations (L24S, del25-27) and G142D in the BA.2 context each partially restored binding, showing the additive effect of antibody escape for the class II, III, and IV mAbs (**Figure 2G**). Surprisingly, reversion of T19I in the BA.2 context failed to restore any binding for mAb CM30, due to the strong escape elicited by the other BA.2 NTD mutations. Thus, BA.1 and BA.2 effectively evade class III and IV mAbs but show different binding of class I and II mAbs. These observations highlight the continued immunological selection to evade potent mAbs elicited by natural infection or immunization^17,44^.

### Mutation context of Omicron RBDs impact mAb escape mechanisms

Despite the significance of immunologic responses to the NTD, mucosal and systemic responses to SARS-CoV-2 infection primarily target the RBD during the acute phase of natural infection^46–48^. Thus, understanding the molecular mechanisms underlying RBD-targeting mAb escape is critical. BA.1 and BA.2 share 12 RBD substitutions, with three additional substitutions (S371L, G446S, and G496S) unique to BA.1 and four substitutions (S371F, T376A, D405N, and R408S) unique to BA.2 (**Figure 3A**). To study the effects of these RBD substitutions on mAb binding we first screened WHU1 spike protein variants containing each of the individual mutations found in BA.1 and BA.2, using the 12 previously described RBD-targeting mAbs (**Figures 3B and S3A**). S2X35, a class I mAb, showed moderately reduced binding due to the E484A mutation. However, each of the S371F, D405N, and R408S AA substitutions caused substantial decreases in binding, likely accounting for the enhanced resistance of BA.2 to mAb S2X35. Class II antibodies C002, C144, and LY-CoV555, and the quaternary binder 2-43, were most affected by the E484A and Q493R single mutations. Class III mAb REGN10987 showed decreased binding with both the N440K and G446S single mutations, and C135 binding was greatly affected by both N440K and Q498R (**Figure 3B**). Two other class III mAbs, C110 and S309, were weakly affected by S317F and were not escaped by the BA.1 and BA.2 spike proteins. Interestingly, S2H97, a class IV mAb, had 4.3-fold decreased binding due to the S371F substitution, versus the 1.4-fold decrease observed with the full set of BA.2 spike mutations. Conversely, no individual AA substitution greatly reduced N3-1 binding despite the strong binding escape observed with the full BA.1 and BA.2 spike proteins.

**Figure 3:**
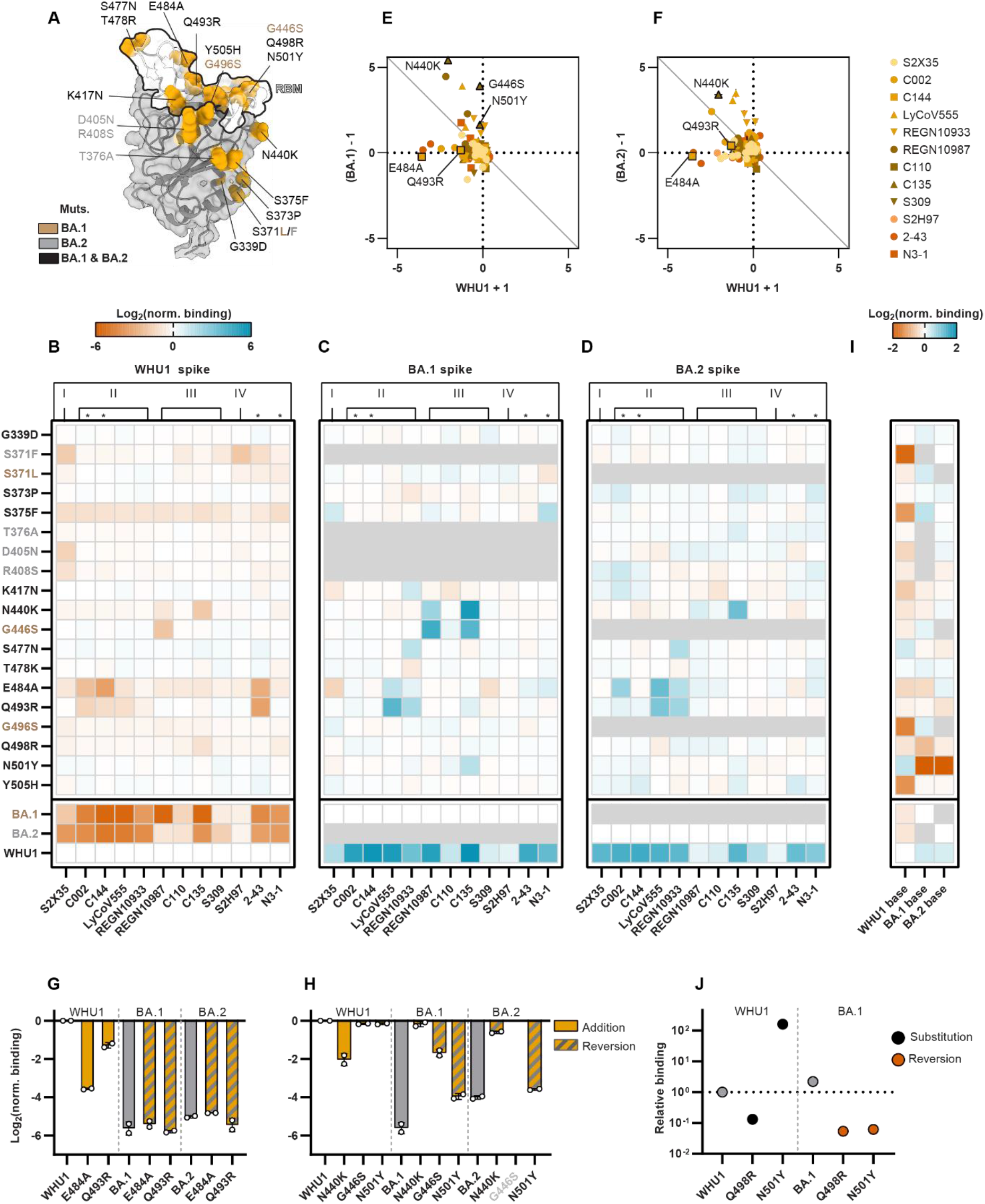
Omicron-RBD mutations enable antibody evasion and preserve hACE2 affinity. A. An enlarged RBD structure (PDB: 7DDN^59^) with mutations in BA.1 (brown), BA.2 (grey), or both (black) indicated. B. Relative monoclonal antibody binding to WHU1 spike proteins containing BA.1- and BA.2-RBD mutations. Red, decreased binding; blue, increased binding; relative to the WHU1 spike. C. Relative monoclonal antibody binding to BA.1 spike proteins containing reversions of the BA.1-RBD mutations to the WHU1 sequence. Colored as in Figure 3B, normalized to the BA.1 spike. D. Relative monoclonal antibody binding to BA.2 spike proteins containing reversions of the BA.2-RBD mutations to the WHU1 sequence. Colored as in Figure 3B, normalized to the BA.2 spike. E. Comparison of the effect of adding (+1) BA.1-RBD mutations to the WHU1 spike protein versus reverting (−1) the corresponding mutations from the BA.1 spike. Mutations with equal antigenic effects in both spike contexts are expected to fall on the diagonal line y = -x. F. Comparison of the effect of adding and reverting BA.2-RBD mutations as in Figure 3E. G. Relative monoclonal antibody C135 binding to the WHU1, BA.1, and BA.2 spike proteins and spike proteins containing the N440K, G446S, or N501Y mutations or reversions as appropriate, normalized to the level of binding to the WHU1 spike. H. Relative monoclonal antibody C144 binding to the WHU1, BA.1, and BA.2 spike proteins and spike proteins containing the E484A or Q493R mutations or reversions, normalized to the level of binding to the WHU1 spike. BA.2 does not contain G446S. I. Relative monomeric hACE2 binding to WHU1 spikes containing BA.1- and BA.2-RBD mutations (WHU1 base), and BA.1 / BA.2 spike proteins containing reversions of the RBS BA.1 / BA.2 mutations (BA.1 base / BA.2 base). Red, decreased binding; blue, increased binding; relative to the WHU1, BA.1, and BA.2 spike proteins as appropriate. J. Monomeric hACE2 binding to WHU1 and BA.1 spike proteins and spike proteins containing the Q498R or N501Y mutations or reversions as appropriate, measured by BLI. All values are normalized to the binding of the WHU1 spike. For all plots, mean ± SD of log-transformed values from at least two biological replicates

Previous studies of Omicron mutations on mAb escape have been performed solely by adding Omicron AA mutations to the WHU1 spike protein^21,22^. These binding or escape measurements fail to capture the nonadditive, epistatic interactions among the mutated sites^49,50^. To explore these contextual effects, we reverted each individual Omicron RBD mutation back to the WT AA in the corresponding BA.1 and BA.2 spike proteins and assayed mAb binding (**Figures 3C-D**). Reversions associated with improved binding, relative to the full set of BA.1 or BA.2 spike mutations, were interpreted to be important for mAb escape in the BA.1 or BA.2 spikes. In the BA.2 spike, no single AA reversion restored mAb S2X35 binding, suggesting an additive effect of S371F, D405N, and R408S mutations together are responsible for escape. LyCoV555 appeared to escape by contributions from E484A and Q493R, which restored binding to different levels in BA.1 and BA.2. REGN10933 binding was greatly restored by both S477N and Q493R reversion in BA.1 and BA.2. However, the K417N reversion substantially restored mAb binding for the BA.1 spike protein but not BA.2. REGN10987 retained affinity to BA.2 due to the absence of the G446S substitution and dampened sensitivity to N440K in the BA.2 context. C135’s escape in BA.2 appears driven by N440K, which completely restored binding when reverted. In BA.1, reverting N440K, G446S, and N501Y all restored binding, suggesting that they all contribute to escape. No reversion in either the BA.1 or BA.2 contexts fully restored binding for the quaternary mAbs C144 or 2-43. Lastly, N3-1 binding improved upon reverting S375F in the BA.1 spike and, to a lesser degree, S373P in BA.2. While these individual mutations do not directly clash with the N3-1 Fabs in the WHU1 structure, there are likely direct or allosteric perturbations to N3-1’s engagement of the Omicron spike (**Figure S3B**), highlighting the importance of mutation context in studying antibody escape pathways.

When comparing the effects of adding and reverting mutations in different spike protein contexts, two interesting cases emerge: (1) mutations that are sufficient for escape in the WHU1 spike but not necessary in the BA.1 or BA.2 spikes, and (2) mutations that are insufficient for escape by the WHU1 spike but necessary for the BA.1 or BA.2 spikes (**Figures 3E-F**). Sufficient but not necessary mutations (below the diagonal line) may indicate examples where BA.1 or BA.2 have stacked multiple mutations that each break a mAb-spike protein binding interaction. For example, C144 binding was strongly disrupted by either E484A or Q493R when these mutations were made to the WHU1 spike but reverting E484A in the BA.1 or BA.2 spike failed to restore binding (**Figure 3G**). Necessary but not sufficient mutations (above the diagonal line) may indicate mutational clusters that have a synergistic or epistatic effect on antibody binding. As an example, N440K reduced, and G446S and N501Y slightly reduced the binding of antibody C135 when made in the WHU1 context (**Figure 3H**). BA.1, which has these three mutations, nearly eliminated C135 binding, and this binding was partially restored when any of the three mutations are reverted. The restoration effect from BA.1 is considerably larger than the reduction in binding from WHU1 suggesting that N440K, G446S, and N501Y have synergistic effects greater than their individual effects. In contrast, BA.2, which lacks G446S, does not show such clear synergy between N440K and N501Y. Overall, these results highlight antigenic differences between the BA.1 and BA.2 spike RBDs and how the overall genetic context impacts antibody escape (**Figure S3C**).

### BA.1 and BA.2 RBDs balance antibody escape and human ACE2 binding

Nearly half of the BA.1 and BA.2 RBD mutations are in the ACE2-binding receptor binding motif (RBM). We screened each single RBD amino acid substitution in the WHU1 spike protein and observed increased hACE2 binding with the S477N and N501Y mutations (**Figure 3I**). Conversely, mutations S371F, S375F, G496S, and Y505H decreased hACE2 affinity. These results are consistent with previously reported RBD DMS measurements^51^, although S371F was more detrimental to hACE2 binding in our assay. We speculate this is due to contextual effects of the full spike glycoprotein.

Reverting each individual RBD mutation in the BA.1 and BA.2 spike proteins showed the deleterious effects of S371F, S375F, K417N, Q496S, and Y505H on hACE2 binding to be less severe in the Omicron contexts. The critical role of N501Y for hACE2 binding by the BA.1 and BA.2 spikes was also shown, as its removal nearly abrogated hACE2 binding, in our assays (**Figures 3I and S3D-E**). Although Q498R mildly reduced hACE2 binding in the WHU1 context, it improved hACE2 binding for the BA.1 spike, and, to a lesser extent the BA.2 spike. Cooperative hACE2 binding due to the Q498R and N501Y substitutions has been previously noted, but to our knowledge the mutation-specific effects had not been measured in the full BA.1 and BA.2 spike proteins^50,52^. We validated these results via biolayer interferometry (BLI). Using dimeric hACE2 we see similar changes in binding (**Figures 3J and S3H**). Together, these results highlight the starkly different molecular basis for hACE2 engagement for BA.1 and BA.2, and the importance of the spike genetic context in understanding these interactions^53–55^.

### Cryptic cross-domain interactions in the BA.1 spike contribute to mAb escape

Reversion of single RBD mutations in the BA.1 and BA.2 spikes broadly failed to fully restore mAbs with quaternary binding modes (C002, C144, 2-43, N3-1) (**Figures S3F-G**). These mAbs simultaneously engage two or three RBDs to enhance their binding via avidity effects. We reasoned that these antibodies can’t bind Omicron relative to the WHU1 spike because of subtle changes in the RBD conformation dynamics. In support of this hypothesis, structures of the BA.1 spike revealed a strict 1-RBD-up, 2-RBD-down conformation^54^. To identify potential cross-domain interactions that may contribute to the extent of escape measured for the full set of BA.1 and BA.2 spike mutations, we created spike proteins containing combinations of NTD, RBD, and S2 mutation sets from the WHU1, BA.1 or BA.2 variants. We then assayed these spike proteins for mAb binding using 12 RBD-targeting mAbs (**Figure 4A**). For most RBD-targeting mAbs, such as C144, the set of BA.1- and BA.2-RBD mutations alone decreased binding to the level of the complete set of BA.1 or BA.2 spike mutations (**Figure 4B**). Interestingly, only the combination of BA.1-RBD and -S2 mutations recapitulate the loss of N3-1 binding measured for the full BA.1 spike (**Figure 4C**). In contrast, the BA.2-RBD mutations alone were adequate for reduced N3-1 binding.

**Figure 4:**
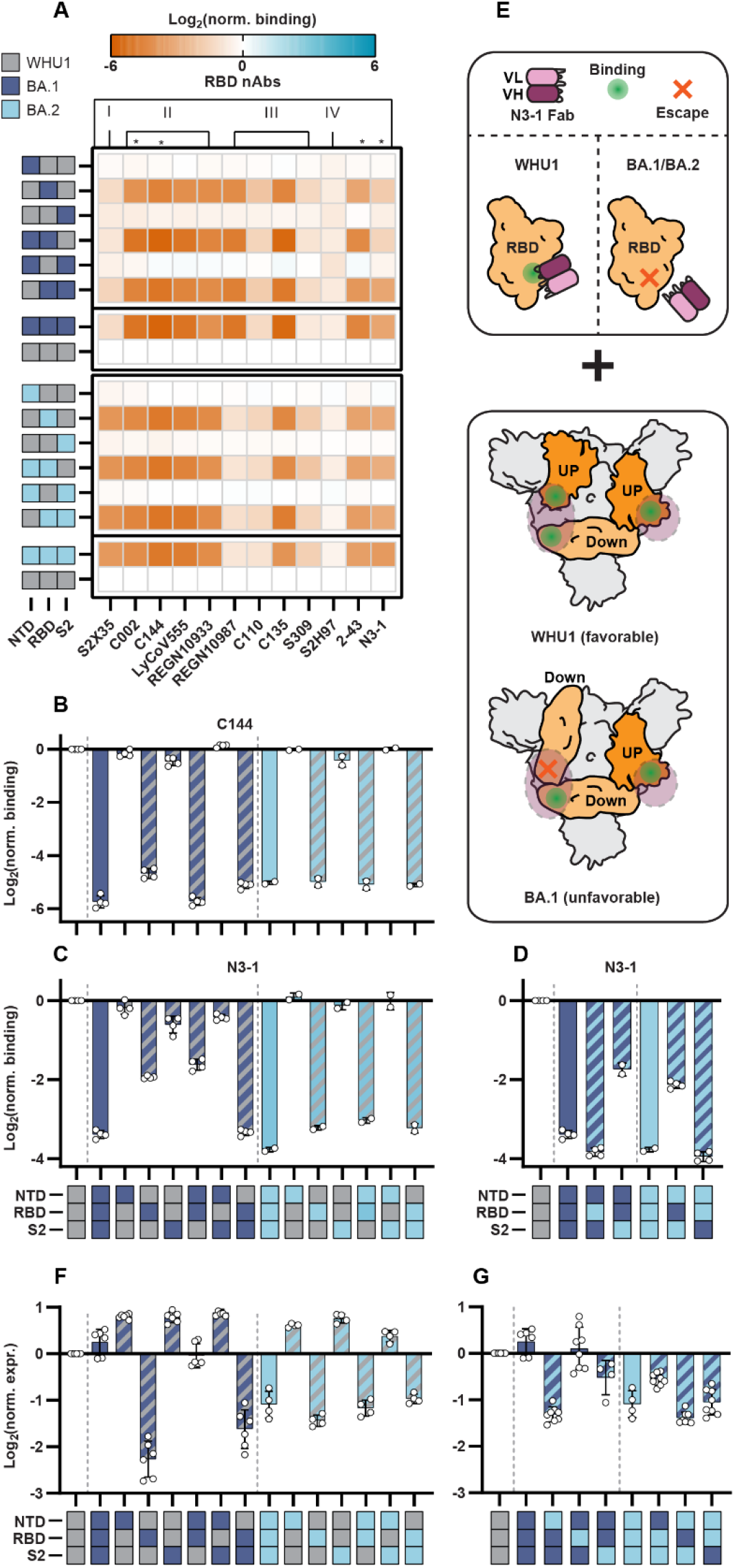
Cross-domain interactions contribute to mAb escape and stabilize the Omicron spike protein. A. Differences in monoclonal antibody binding of RBD-directed antibodies against spike proteins containing combinations of the NTD, RBD, and S2 mutation sets from the BA.1 and BA.2 variants. Red, decreased binding; blue, increased binding, relative to the WHU1 spike. B. Monoclonal antibody binding of C144 to spike proteins containing combinations of the NTD, RBD, and S2 BA.1 and BA.2 mutation sets relative to WHU1. C. Monoclonal antibody binding of N3-1 to spike proteins containing combinations of the NTD, RBD, and S2 BA.1 and BA.2 mutation sets relative to WHU1. D. Additional N3-1 binding data as in Figure 4C. E. Proposed escape mechanism for biparatropic antibody (N3-1) by Omicron spike proteins. Antibody binding reduced via mutations at the N3-1 binding epitope on the RBD (top). Top-down view of spike protein trimers with annotated RBD up vs down positions (bottom). Violet circles, N3-1 binding epitope; green dot, preserved binding; red “x,” escaped binding. F. Differences in expression of spike proteins containing combinations of the NTD, RBD, and S2 BA.1 and BA.2 mutation sets. Data normalized to WHU1 spike expression. G. Additional expression data as in Figure 4F. For all plots, mean ± SD of log-transformed values from at least two biological replicates

To determine if differences in the ability of BA.1 and BA.2 to escape of N3-1 was intrinsic to their RBD and S2 mutations, we created chimeric spikes by swapping the BA.1 and BA.2 mutation sets (NTD, RBD, and S2) (**Figure 4D**). We observed a further 1.4-fold reduction in N3-1 binding when the BA.1-RBD was replaced with the BA.2-RBD, in the BA.1 spike. Furthermore, the BA.2-S2 mutations did not synergistically reduce N3-1 binding when combined with the BA.1-NTD and -RBD mutations. Conversely, replacing BA.2-RBD mutations with BA.1-RBD mutations dampened N3-1 escape. By swapping the BA.1-S2 mutations into the BA.2 spike, we did observe a marginal (1.1-fold) reduction in N3-1 binding. We postulate that the N856K and L981F mutations, which comprise the only differences between the BA.1- and BA.2-S2 mutations sets, alter the RBD-up vs -down equilibrium or spike conformation, thus further reducing N3-1 binding. These data highlight possible routes of quaternary-binding mAb evasion by Omicron through altering antigenic epitopes and RBD dynamics (**Figure 4E**).

### Omicron spike domains provide stability compensation to immune evasive RBD

Expression-–a proxy for spike stability—correlates with improved infectivity and viral fitness^56,57^. BA.1 spikes express 1.2-fold greater than WHU1 spikes while BA.2 express 2.1-fold lower (**Figure S1D**). We determined the combination of mutations that are responsible for this increased expression by establishing the effect of each individual BA.1 and BA.2 mutation. Spike expression was monitored via fluorescent signals from two discrete epitopes: a triple FLAG in the linker between the ectodomain and transmembrane domain and the foldon trimerization domain (**Figure S4A-B**) (See Methods). Most of the BA.1 and BA.2, NTD and S2 mutations enhanced spike expression. For example, NTD mutations del69-70 and ins214EPE in BA.1, and G142D shared by both BA.1 and BA.2 improved WHU1 spike expression. Reversion of either del69-70 or ins214EPE in the BA.1 spike only modestly decreased expression (**Figure S4C**). Interestingly, the G142D mutation is more central to the overall stability of the BA.2 spike, as its reversion reduced BA.2 spike expression 5.3-fold. Surprisingly, we also saw extremely destabilizing mutations in the BA.1 and BA.2 RBDs. The addition of S375F to the WHU1 spike dramatically reduced expression 12.3-fold. Mutations N440K and E484A, which are responsible for mAb escape, and N501Y, which enhances hACE2 affinity are also mildly destabilizing (**Figure S4D**).

Next, we assayed each set of domain-specific mutations (NTD, RBD, and S2) in the context of WHU1 spike (**Figure 4F**) to determine their relative effects on spike expression. We found that both the BA.1- and BA.2-RBD mutation sets greatly reduced spike protein expression relative to WHU1, whereas the NTD and S2 mutation sets increased expression. The fully mutated BA.1 spike had greater expression than the WHU1 spike, suggesting that the destabilizing mutations in the RBD are compensated for primarily by mutations in the NTD and S2 domains. Further domain exchanges showed that the BA.1 NTD mutations are sufficient to stabilize the BA.1 RBD mutations, with the S2 mutations contributing, but insufficient on their own. BA.2 follows the same general pattern, however the BA.2-NTD and -S2 were less effective at offsetting the relatively milder decreased expression associated with the BA.2-RBD. This finding is consistent with the relatively poor spike expression in BA.2, compared to expression by the WHU1 and BA.1 variants. Lastly, we swapped BA.1 and BA.2 domains for the BA.1 and BA.2 spikes to determine if stabilizing effects were transferrable. Interestingly, the exchange of BA.2 NTD mutations into the BA.1 spike greatly reduce spike expression (**Figure 4G**). The exchange of BA.1 RBD mutations into the BA.2 spike also reduced BA.2 spike expression.

To test how expression relates to spike stability, we tested the thermal denaturation of a subset of spike variants via differential scanning fluorimetry (DSF) (see Methods). Soluble SARS-CoV-2 spike trimers generate two distinct denaturation peaks, denoted here as Tm1 and Tm2. Compared to other VOCs, the BA.1 and BA.2 spike proteins have poor thermostability, as shown by their respective 7 °C and 6 °C shifts in Tm1, relative to WHU1 (**Figures S4A-B**). DSF measurements of spike variants reveals that these effects are driven by RBD mutations in BA.1 (**Figures S4C-F**) and BA.2 (**Figures S5G-J**). In sum, both BA.1 and BA.2 spikes are destabilized by their highly mutated RBDs. Although the BA.1 NTD mutations don’t improve the thermostability of the spike protein, they compensate for poor spike expression. Taken together, these results reveal that the expression loss due to nAb-evasive but otherwise destabilizing RBD mutations is offset by otherwise stabilizing NTD & S2 mutations. These mutations work synergistically with a particular spike background, as swapping mutation sets between BA.1 and BA.2 leads to overall reduced expression.

## Discussion

This work dissects the effects of individual mutations in different spike protein contexts to understand how these mutations evade neutralizing antibodies and impact spike stability. The G142D, del143-145 mutation cluster in the BA.1-NTD is both necessary and sufficient for the comprehensive escape of class I, III, and IV NTD-targeting mAbs (**Figure 2F**). All BA.2-NTD mutations, except for V213G, collectively contribute to the escape of class II, III, and IV NTD-targeting mAbs (**Figure 2G**). These results highlight antigenic differences between the BA variants and the original variant, as well as between the BA.1 and BA.2 spike NTDs themselves. Moreover, our work reveals that multiple NTD mutations additively evade potent NTD-directed mAbs.

In contrast to the NTD, mutations in the BA.1- and BA.2-RBDs often resulted in non-additive levels of escape from RBD-targeting mAbs. We identified instances where multiple mutations were required to escape binding completely. For example, reversions of either N440K or G446S in the BA.1 spike largely restored the binding of REGN10933 and C135 (**Figure 3C**). We also identified several class II mAbs (C002 and C144) that failed to show binding improvements after single mutation reversions in the Omicron spike proteins, suggesting redundant mutations contributing to antigenic escape (**Figure 3B-D**). We speculate that Omicron RBDs have undergone extensive mutation under continuous pressure to evade diverse classes of RBD-targeting antibodies, outside of the predominant class II antibodies found in polyclonal plasmas after immunization or natural infection^32^. This redundancy in escape may also arise from mutation-induced alterations in RBD conformational equilibria and dynamics, as described in previous structure studies^53,54,58^.

Reverting individual BA.1 and BA.2 mutations back to WT (WHU1) in the BA.1 and BA.2 spikes generally failed to restore binding for the four mAbs we tested that were capable of binding multiple RBDs at once: C002, C144, 2-43, and N3-1 (**Figures S3F-G**). These results suggested that the virus was likely not simply presenting different epitopes than the ancestral variant but presenting them in the context of different proportions of RBD up vs down states. We showed that RBD-S2 cross-domain interactions in the BA.1 spike led to reduced N3-1 binding beyond the BA.1-RBD mutations alone (**Figure 4C**). In further support of this hypothesis, the removal of the N856K and L981F mutations from the BA.1 spike, which are implicated in creating cross-domain interactions, partially restored N3-1 binding to the level of the BA.1-RBD mutations alone (**Figure 4D**). Similarly, BA.2-RBD mutations proved even more effective than BA.1-RBD mutations for escaping N3-1, in line with recently solved structures that show that the down conformation of the RBD is largely stabilized in the BA.2-RBD mutations^58^.

The accumulation of novel mutation clusters in Omicron, likely due to selection pressure to evade immunity, came at the cost of destabilizing the RBD. Several RBD mutations, most notably S375F, drastically reduce spike expression (**Figure S4D**). We propose that NTD mutations such as the 69-70 deletion offset protein folding/stability deficiencies associated with Omicron RBDs^57^. Addition of BA.1-NTD mutations compensated for the poor expression of the BA.1-RBD mutations (**Figure 4F**). Interestingly, the same compensation effects were not seen with the BA.2-NTD and BA.2-RBD mutations, resulting a lower net expression of BA.2 spike relative to WHU1 in our assays. Future studies will be required to explore whether and how improvements in immune evasion have led to fitness costs for Omicron relative to spike expression and fitness.

Our data suggest that the Omicron BA.1 and BA.2 subvariants retain high affinity for hACE2. Previous directed evolution^52^ and YSD-DMS^50^ studies have shown N501Y to greatly improve hACE2 binding and Q498R to moderately reduce it; these studies also determined that the co-occurrence of N501Y and Q498R synergistically boosted hACE2 affinity. We confirm that the full Omicron BA.1 and BA.2 spikes also benefit from these substitutions, with N501Y and Q498R playing critical roles in the molecular engagement of hACE2 (**Figure 3I-J**). Intriguingly, this effect appears distinct in BA.1 versus BA.2 (**Figure S3D-E**). As compared to BA.1, N501Y has a much greater impact on ACE2 binding in BA.2 but Q498R is not as critical, possibly due to the absence of BA.1 mutations G446S and G496S. Taken together, these results reveal evolutionary features of Omicron spikes that enable the accruement of immune evasive mutations without sacrificing hACE2 affinity and infectivity.

Our study has several limitations. First, we used a prefusion stabilized spike protein that does not precisely mimic the dynamics of the native Omicron spike protein^26^. Second, our binding assays use a set of potent neutralizing mAbs which only serve as proxies for the antibodies found in patient antibody repertoires after immunization or natural infection. Third, our work only touches on antibody recognition and hACE2 binding; T-cell immunity plays a critical role in protecting against SARS-CoV-2 disease. Additional studies focused on the perturbations of spike variants on T-cell response will continue to bridge the gap in the understanding of immune escape between humoral and cell-mediated immunity.

In the aggregate, the data presented here add critical new information about key features of Omicron spike protein mutations and how these mutations synergize to create spike variants that successfully evade antibodies while maintaining high affinity hACE2 binding. Our binding maps largely complement prior structure-based studies of binding escape, but now provide new insights into the role of compensatory substitutions in the NTD that impact both expression/stability and conformation. We conclude that the continuing accumulation of NTD mutations will further alter the conformational equilibrium and stability of the spike protein to allow for the accumulation of new, more virulent mutations in the RBD. Further, our study also highlights the importance of rapidly analyzing novel variant spike proteins in near-native contexts. As SARS-CoV-2 continues to evolve and new variants inevitably arise and spread, it is critical that these mutations can be understood in their native genetic contexts to better inform future therapeutic targets and vaccine formulations. Finally, the strategy we have used here will be applicable to future zoonotic outbreaks and other heretofore unrelated microbial pathogens. Mammalian cell-display will continue to serve as a powerful platform for investigating evolutionary trajectories of infectious agents and engineering conformational vaccine candidates.

## Acknowledgments

We thank Drs. Marc Boom and Dirk Sostman for ongoing support; and Dr. Sasha M. Pejerrey, Dr. Heather McConnell, and Ms. Adrienne Winston for editorial contributions. The research was supported in part by the Houston Methodist Academic Institute Infectious Diseases Fund and many generous Houston philanthropists (JMM and JDG). We are especially grateful to Carole Walter Looke and Jim Looke for their generous philanthropic gift to ADAPT (JDG). The funders had no role in the design and conduct of the study; collection, management, analysis, and interpretation of the data; preparation, review, or approval of the manuscript; and decision to submit the manuscript for publication. KJ was supported by the Provost’s Graduate Excellence Fellowship (PGEF) at UT Austin. The research was also supported by a Cooperative Agreement (W911NF-17-2-0091) between ARL and UT Austin to ADE and JDG, the Bill and Melinda Gates Foundation (KJ, CWC, AA, and IJF), and the Welch Foundation (F-1808 to IJF).

## Author Contributions

KJ, THSS, IJF, and JDG designed the research. KJ performed the flow experiments. KJ, DRB, AA, and JDG purified antibodies and other reagents. KJ and THSS cloned spike variants. KJ and CWC performed BLI DSF experiments. RJO and JMM isolated and sequenced Omicron. XX, HX, PS, and SW provided the authentic virus neutralization data. KJ, THSS, CDJ, and JDG analyzed the data. KJ, THSS, ADE, IJF, and JDG wrote the paper with editorial assistance from all co-authors.

## Inclusion and Diversity

One or more of the authors of this paper self-identifies as an underrepresented ethnic minority in science. The author list of this paper includes contributors from the location where the research was conducted who participated in the data collection, design, analysis, and/or interpretation of the work.

## MATERIALS & METHODS DETAILS

### Spike variant cloning

We assembled protein variants using a Golden Gate / TypeIIs cloning framework. Generally, segments of the spike protein coding sequence were ordered as synthetic DNA fragments (IDT, eBlocks), cloned into a plasmid backbone, and sequence verified. Verified DNA parts were transferred with plasmid backbone into 96-well PCR plates using an Echo 525 liquid handler (Beckman Coulter). To each well, we added the following Golden Gate reaction mixture: 0.25 μL of T4 or T7 DNA Ligase (NEB, M0202 / M0318), 0.25 μL of AarI (Thermo Fisher, ER1582), 0.2 μL AarI Oligo (Thermo Fisher), 1 μL T4 DNA Ligase Buffer (NEB B0202A), and nuclease-free water to bring the final volume to 10 μL per reaction. We incubated the reaction mixtures on a thermocycler using the following settings: 25 digestion and ligation cycles (1 min at 37°C and 2 min at 16°C), a final digestion step (30 min at 37°C), and heat inactivation (20 min at 80°C). For assemblies with 4+ parts, we increased the cycled digestion and ligation steps to 3 and 5 min, respectively, to improve assembly efficiencies.

We transformed each unique reaction mixture into NEB10-beta cells (NEB, C3019) using the Mix & Go! E.coli Transformation Buffer Set (Zymo Research, T3002). Cell were outgrown with SOC at 37°C and drop-plated onto LB agar + carbenicillin in Nunc OmniTrays (5 μL per spot, 96-spots per plate) (Thermo Fisher, 140156). We allowed the drops to dry at room temperature before transferring the plates to an incubator at 37°C for growth overnight. After growth, we picked colonies into deep well grow blocks (Axygen P-2ML-SQ-C-S or Greiner Bio-One, 780270) containing 1 mL of Superior Broth (AthenaES, 0105) media + carbenicillin and grew them overnight at 37°C with shaking. Overnights were miniprepped using the Wizard SV 96 Plasmid DNA Purification Kit (Promega A2250) and sequence-verified before further characterization.

### Expression and purification of neutralizing anti-spike monoclonal antibodies

We cultured Expi293 cells in Expi293 Expression Medium (Sigma-Aldrich A1435101) and used a humidified cell culture incubator to maintain cells at 37°C and 8% CO_2_ with continuous shaking at 125 rpm. For transfection, we used an Expi293 Transfection Kit (Sigma-Aldrich L3287) according to the manufacturer’s instructions. Briefly, we transfected cells with VH and VL expression vectors at a 1:3 molar ratio. Five days after transfection, we collected the protein-containing supernatant using a two-step centrifugation protocol. First, we separated cells and supernatant by centrifuging cultures at 4°C and 300 g for 5 min. Next, we separated cell debris and supernatant by centrifuging at 4°C and 3,000 g for 25 min. To purify human IgGs, we washed Protein G magnetic beads (Promega G7471) with PBS buffer and added the beads to the separated supernatant in a 1:10 volumetric ratio. After a 1h incubation with gyration at room temperature, we used a magnetic peg stand to pellet bead-bound antibodies, which we washed before final elution with a 100 mM glycine-HCl solution at pH 2.5. Finally, we passed the elute through a 0.22-μm syringe filter to clarify residual beads before neutralization with 2 M Tris buffer at pH 7.5. We kept purified antibodies at 4°C or −20°C for short and long-term storage, respectively.

### Expression and purification of human ACE2-Fc

We recombinantly expressed human ACE2-Fc in Expi293 cells using a previously described method with minor modification^62^. Briefly, we transfected the ACE2-Fc expression vector into Expi293T cells (Sigma-Aldrich). Five days after transfection, we centrifuged the cultures at 4°C and 300 g for 5 min and collected the supernatant. We further separated cell debris and supernatant by centrifuging at 10,000 g and 4°C for 20 min. After resuspending it in PBS, we purified ACE2-Fc over Protein A Agarose (Thermo Fisher 15918014). We next equilibrated the Protein A Agarose in PBS buffer, ran through the supernatant three times, and used 10 bed volumes of PBS buffer for washing. Finally, we used 100 mM glycine-HCL at pH 2.4 to elute ACE2-Fc into 0.1× volume Tris buffer at pH 8.5 and 100 mM NaCl. We kept purified ACE2-Fc at 4°C and −20°C for short and long-term storage, respectively.

### Expression and purification of SARS-CoV-2 spike proteins

We transfected and expressed plasmids using Expi293 cells using previously described methods^26^. Briefly, we purified variants from 40 mL of cell culture. We filtered the supernatant using a 0.22-μm filter and ran it through a StrepTactin Superflow column (IBA 2-1206-025). We further purified spikes via Superose 6 Increase 10/300 (GE GE29-0915-96) size-exclusion column chromatography with a buffer containing 2 mM Tris at pH 8.0, 200 mM NaCl, and 0.02% NaN3. We kept purified samples at 4°C and − 20°C for short and long-term storage, respectively.

### HEK293T transfection

We seeded cells into 6 or 12-well polystyrene-coated plates (VWR 10861-696, 10861-698) at a density of 0.3 × 10^6^ cells mL^−1^ or 0.1 × 10^6^ cells mL^−1^, respectively, one day before transfection. At 60-80% confluence, we used Lipofectamine 3000 (Thermo Fisher L3000015) and Opti-MEM (Gibco 51985091) to transfect cells with expression plasmids (3 μL of lipofectamine per μg of plasmid DNA) according to manufacturer instructions. At 48 h post-transfection, we assayed or collected the cells.

### Flow cytometry and data analysis

At 48 h post-transfection, we collected HEK293T cells containing surface-displayed spike. We washed cells once with PBS and used gentle pipetting to resuspend them in PBS. We used the Logos Biosystems (L40002) cell counter to determine cell density and spun cells down at 200 g for 1 min. We next decanted the supernatant and resuspended cells to a density of ~3 × 10^6^ cells mL^−1^ in chilled PBS-BSA using 1% BSA (Sigma-Aldrich A3294), 1X PBS, and 2 mM EDTA at pH 7.4.

We used Axygen deep well grow blocks (P-2ML-SQ-C-S) to prepare flow cytometric assays. We added predetermined concentrations of primary antibody or chimeric cell receptor (ACE2-Fc) diluted in PBS-BSA and 50 μL (1.5 × 10^5^) of HEK293T cells to each well. We incubated the mixtures at room temperature for 1 h with shaking at 950 rpm. To pellet cells, we spun plates at 500 g in a swinging bucket rotor for 2 min. We then washed cells twice by decanting the supernatant and adding 500 μL of PBS-BSA to each well. To each well, we added 500 μL total volume of 5 μM Alexa Fluor® 488 anti-mouse secondary (SouthernBiotech 1031-30) and 10 μM Alexa Fluor® 647 anti-human secondary (SouthernBiotech 2048-31) antibodies in PBS-BSA. We incubated plates in the dark for 25 min at 4°C with shaking at 950 g. We washed each well twice with PBS-BSA and resuspended cells in PBS-BSA (300 μL) before loading them onto the SA3800 Spectral Cell Analyzer (SONY).

To establish forward scatter-area (FSC-A) and side scatter-area (SSC-A) gating, we used HEK293T cells. For singlet discrimination, we gated with forward scatter-height (FSC-H) vs forward scatter-area (FSC-A) and side scatter-height (SSC-H) vs side scatter-area (SSC-A). For each assayed sample, we acquired a minimum of 10,000 singlet events. We further analyzed singlet HEK293T cells using Alexa Fluor 488 (AF-488) and Alexa Fluor 647 (AF-647) channels with excitation and detection settings recommended by the manufacturer. To reduce spectral spill-over and autofluorescence effects, we applied spectral unmixing to all data.

For each sample, we measured the median height (*H*) for the AF-488 and AF-647 channels. To measure spike expression, we used the signal from the AF-488 channel (anti-FLAG). We used the following equation to calculate the expression of spike variant (*x*) relative to WT (*6P-D614G*):

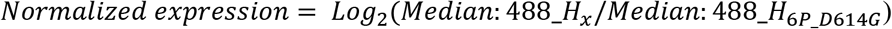

To correct for changes in transfection efficiency or spike expression in antibody or ACE2 binding measurements, we also included anti-FLAG signal as an internal normalization control. We used the following equation to calculate normalized binding measurements of spike variant (*x*) expression relative to WT (*6P-D614G*):

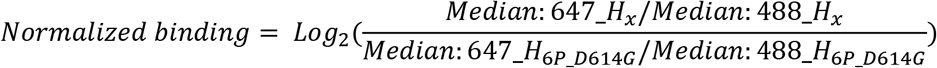

We used FlowJo v9 for all flow cytometry data analyses.

### SARS-CoV-2 authentic virus neutralization assay

To measure human sera and monoclonal antibody neutralization titers, we used a fluorescent focus reduction neutralization test (FFRNT) with an mNeonGreen (mNG) reporter SARS-CoV-2 (strain USA-WA1-2020) or SARS-CoV-2 (strain USA-WA1-2020) bearing a variant spike gene (e.g., Omicron BA.1). The construction of the mNG USA-WA1-2020 SARS-CoV-2 bearing variant spikes has been reported previously (Liu et al., 2021; Zou et al., 2022). For the FFRNT assay, we seeded 2.5 × 10^4^ Vero E6 cells into black, μCLEAR flat-bottom 96-well plates (Greiner Bio-One™). We incubated plates at 37°C with 5% CO_2_ overnight. The next day, each sample was two-fold serially diluted in culture medium with an initial dilution of 1:20. We incubated diluted serum or antibody with 100-150 fluorescent focus units (FFU) of mNG SARS-CoV-2 at 37°C for 1 h before loading the serum-virus mixtures into 96-well plates pre-seeded with Vero E6 cells. Following a 1 h infection period, we removed the inoculum and added overlay medium (100 μL DMEM + 0.8% methylcellulose, 2% FBS, and 1% penicillin/streptomycin). We then incubated the plates at 37°C for 16 h and acquired raw images of mNG fluorescent foci using a Cytation™ 7 (BioTek) cell imaging reader with a 2.5× FL Zeiss objective and wide field of view. We used GFP software settings [469,525], a threshold of 4000, and an object selection size of 50-1000 μm during image processing. For relative infectivity calculations, we counted and normalized the foci in each well relative to non-serum/antibody-treated controls. We plotted curves of relative infectivity versus serum dilution using Prism 9 (GraphPad). We used a nonlinear regression method to determine the dilution fold at which 50% of mNG SARS-CoV-2 was neutralized, defined as FFRNT50 in GraphPad Prism 9. Each antibody was tested in duplicate.

### Biolayer Interferometry

After 3C protease-mediated cleavage, we diluted supernatants containing spike variants two-fold with BLI Kinetics Buffer (Fortébio) containing 10 mM HEPES at pH 7.5, 150 mM NaCl, 3 mM EDTA, 0.05% v/v Surfactant P20 (Cytiva BR100054), and 1 mg mL^−1^ BSA. We also serially diluted analytes with the BLI buffer. We hydrated anti-mouse Fc capture (AMC) biosensors (FortéBio 18-5088) in BLI buffer for 10 min in an Octet RED96e (FortéBio) system and then immobilized mouse anti-FLAG M2 (Sigma-Aldrich F3165) antibodies to the AMC sensor tips. For each assay, we performed the following steps: 1) baseline: 60 s with BLI buffer; 2) IgG immobilization: 360 s with anti-FLAG IgG; 3) spike loading: 360 s with diluted supernatants; 4) baseline: 300 s with BLI buffer; 5) association: 600 s with serially diluted analytes (antibodies or ACE2); 6) dissociation: 600 s with BLI buffer. We used Octet Data Analysis software v11.1 with steady-state analysis to reference-subtract and analyze the data.

### Differential Scanning Fluorimetry

Sample solutions containing 5X SYPRO Orange Protein Gel Stain (Invitrogen S6651) and 0.15-0.20 mg/mL of purified spike protein were added to a 96-well qPCR plate (Azenta Life Sciences 4ti-0955). Fluorescence measurements were obtained continuously using λex=465 nm and λem=580 nm, using a Roche LightCycler 480 II, and a temperature ramp rate of 4.5°C/minute increasing from 22 °C to 95 °C. Fluorescence data were then plotted as the derivative of the melting curve as a function of temperature (-dF/dT). SARS-CoV-2 spike proteins generate two local minimums that we report to as Tm1 and Tm2. All data were visualized in GraphPad Prism 9. For simplicity, we normalize all melting curve traces by their absolute min -df/dT value.

### Computational analysis of GISAID sequence data

To investigate the clinical frequency of SARS-CoV-2 spike mutations and the probability of mutation co-occurrences, we performed pairwise amino acid sequence alignments between the GISAID spike reference sequence, GenBank number QHR63250.2, and all GISAID EpiCoV database SARS-CoV-2 spike amino acid sequences. We downloaded amino acid sequences from GISAID (accessed on December 18, 2021) as a FASTA file. We performed semi-global amino acid sequence alignment using MATLAB’s Needleman-Wunsch alignment function as a part of its Bioinformatics Toolbox add-on. We set alignment parameters to include no gap open penalty at the beginning and end of sequences, an internal gap open penalty of 5, a gap extension penalty of 2, and the BLOSUM80 scoring matrix, as the aligned sequences were similar.

We filtered alignment pairs to remove all sequences that were non-human in origin, sequences containing over 1280 or fewer than 1260 amino acids, and sequences containing more than 800 unknown (“X”) amino acids. We identified non-synonymous mutations from the alignments. The following equation was used to find frequencies for each mutation:

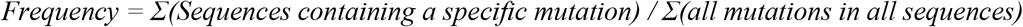

We found mutations that occurred independently by removing all alignment pairs that did not contain the target mutation and sequences that contained mutations other than the target mutation. Additionally, we considered alignment pairs for which the only other mutation (other than the target) was D614G were as independent, as D614G was highly prevalent in all strains after its initial appearance. The following equation was used to calculate frequency for independent mutations:

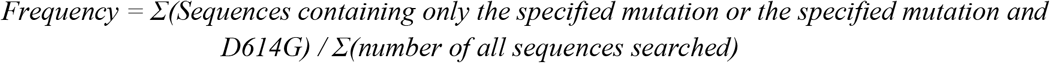

### Structural analyses and data visualization

We downloaded all structures (7DDN^61^, 7L2D^29^, and 7L2E^29^) as PDB files from the RCSB PDB and imported them into ChimeraX 1.1. We rescaled the Log2 (normalized binding) values (−7 to 0) and converted them to monochromatic ChimeraX color codes representing changes in binding relative to 6P-D614G spike. Dark red indicates decreased binding and white indicates no change in binding. For every amino acid screened in our Spike Display assay, we superimposed these colors scales onto spike protein structures. In figures showing grouped antibody epitopes, we averaged normalized binding values for all mutations in each position for every antibody comprising that group. Finally, we converted the averaged binding values to a single color that was then mapped onto spike structures.

## STATISTICAL ANALYSIS

The means ± S.D. log-transformed values, or non-transformed values were calculated and reported for all appropriate data.

