## Supplementary information for "Antibody escape and cryptic cross-domain stabilization in the SARS-CoV-2 Omicron spike protein"

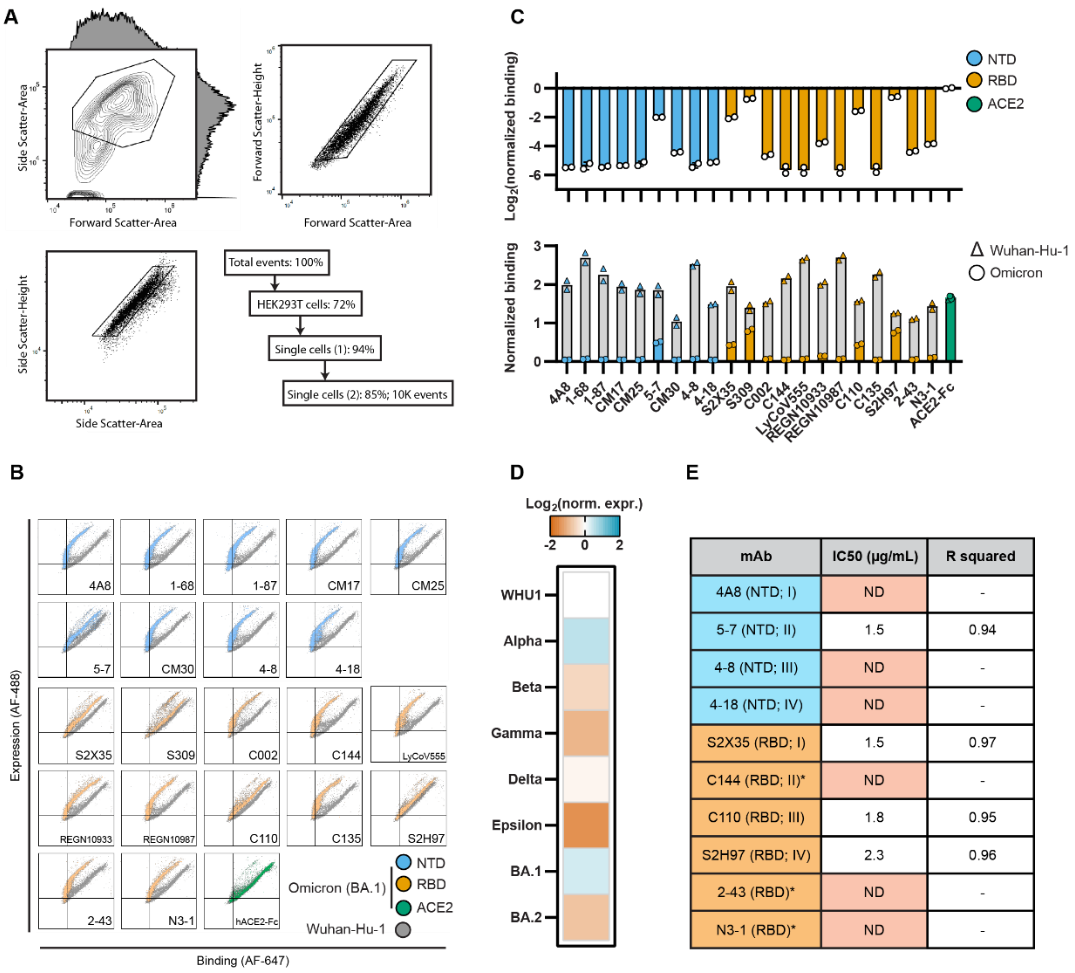


| mAb | IC50 (μg/mL) | R squared |
|---|---|---|
| 4A8 (NTD; I) | ND | - |
| 5-7 (NTD; II) | 1.5 | 0.94 |
| 4-8 (NTD; III) | ND | - |
| 4-18 (NTD; IV) | ND | - |
| S2X35 (RBD; I) | 1.5 | 0.97 |
| C144 (RBD; II)* | ND | - |
| C110 (RBD; III) | 1.8 | 0.95 |
| S2H97 (RBD; IV) | 2.3 | 0.96 |
| 2-43 (RBD)* | ND | - |
| N3-1 (RBD)* | ND | - |

**Figure S1**: Spike Display enables rapid characterization of spike proteins variants

A. Flow cytometry gating (shown with data for BA.1 in Figure 1F; see Methods).
B. Representative flow cytometry data for WHU1 spike protein (grey) and BA.1 spike protein (colors) labeled for expression (y-axis) and antibody / ACE2 binding (x-axis). A shift to the left between WHU1 and BA.1 shows reduced antibody / ACE2 binding at the comparable levels of spike protein expression. Median fluorescence values used to calculate normalized binding for each spike protein variant and antibody / ACE2 combination (see Methods).
C. Quantitative data for the BA.1 and WHU1 rows in Figure 1F. Normalized change in binding (top) shows the difference in normalized monoclonal antibody and ACE2 binding between BA.1 and WHU1. Normalized binding (bottom) shows the binding values for these antibodies and ACE2 for both WHU1 (grey) and BA.1 (colors) relative to expression. Bar graphs represent mean ± SD of log-transformed values from at least two biological replicates.
D. Relative expression values for the spike protein variants shown in Figure 1F, normalized to WHU1 expression.
E. IC50 and fit data calculated from the plots shown in Figures 1G and 1H.

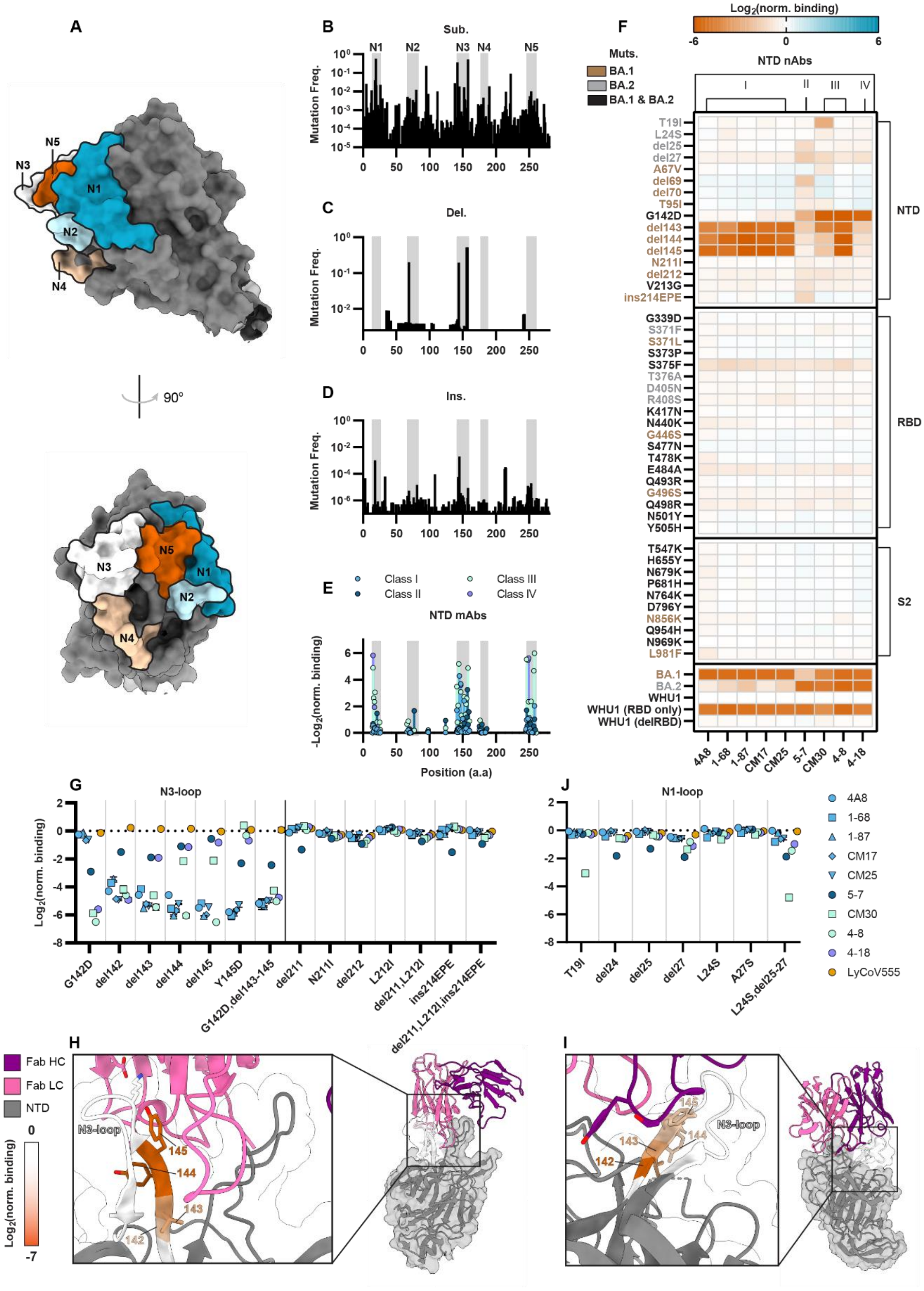

A
N1
N2
N3
N4
N5
90°
B
Sub.
C
Del.
D
Ins.
Mutation Freq.
E
Class I
Class II
Class III
Class IV
NTD mAbs
-Log2(norm. binding)
Position (a.a)
F
Log2(norm. binding)
Muts.
BA.1
BA.2
BA.1 & BA.2
NTD nAbs
I
II
III
IV
T19I
L24S
del25
del27
A67V
del69
del70
T95I
G142D
del143
del144
del145
N211I
del212
V213G
ins214EPE
G339D
S371F
S371L
S373P
S375F
T376A
D405N
R408S
K417N
N440K
G446S
S477N
T478K
E484A
Q493R
G496S
Q498R
N501Y
Y505H
T547K
H655Y
N679K
P681H
N764K
D796Y
N856K
Q954H
N969K
L981F
BA.1
BA.2
WHU1
WHU1 (RBD only)
WHU1 (delRBD)
NTD
RBD
S2
4A8
1-68
1-87
CM17
CM25
5-7
CM30
4-8
4-18
G
N3-loop
Log2(norm. binding)
G142D
del142
del143
del144
del145
Y145D
G142D,del143-145
del211
N211I
del212
L212I
del211,L212I
ins214EPE
del211,L212I,ins214EPE
J
N1-loop
T19I
del24
del25
del27
L24S
A27S
L24S,del25-27
4A8
1-68
1-87
CM17
CM25
5-7
CM30
4-8
4-18
LyCoV555
H
Fab HC
Fab LC
NTD
Log2(norm. binding)
N3-loop
142
143
144
145
I
N3-loop
142
143
144
145

**Figure S2**: Immunologic selection to evade mAbs at the NTD supersite.

A. NTD domain structure (PDB: 7DDN[61]) with colored and labeled N-Loops (N1-N5).
B. Frequency of NTD-substitutions.
C. Frequency of NTD-deletions.
D. Frequency of NTD-insertions (GISAID accessed on 18/December/2021; see Methods).
E. Spike display binding measurements of single alanine substitutions at each N-loop position. Dots represent mean binding values for each class of mAbs. These data were previously published in Javanmardi *et al*. 2021 and reanalyzed for this figure[23].
F. Binding of NTD-targeting mAbs to WHU1 spikes containing BA.1- and BA.2- mutations normalized to WHU1 spike. Red: decreased binding; blue: increased binding.
G. Differences in normalized binding of monoclonal antibodies caused by mutations in the N3-loop and the set of 211-214 mutations found in BA.1. Data represent mean ± SD of log-transformed values from at least two biological replicates.
H. Structure (PDB: 7L2D[29]) of 1-87 Fab (light chain, pink; heavy chain, violet) in complex with NTD (gray) with highlighted N3-loop (white). Binding data from 1-87 in Fig. S2H is superimposed on the N3-loop.
I. Structure (PDB: 7L2E[29]) of 4-18 Fab (light chain, pink; heavy chain, violet) in complex with NTD (gray) with highlighted N3-loop (white). Binding data from 4-18 in Fig. S2H is superimposed on the N3-loop.
J. Differences in normalized binding of monoclonal antibodies caused by mutations in the N1-loop mutations found in BA.2. Data are a mean ± SD of log-transformed values from at least two biological replicates.

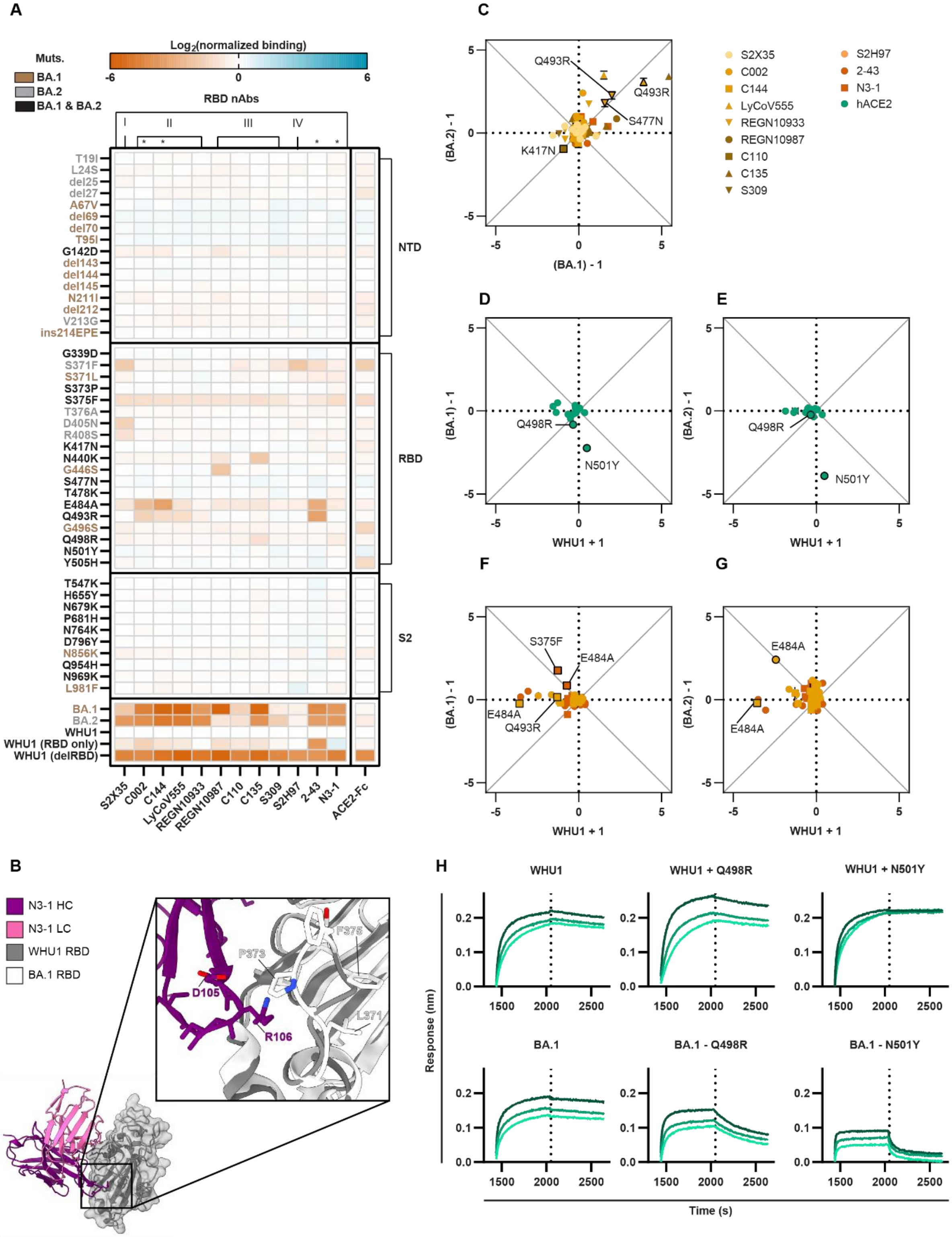
A
Muts.
BA.1
BA.2
BA.1 & BA.2
Log2(normalized binding)
-6
0
6
RBD nAbs
I
II
III
IV
NTD
RBD
S2
T19I
L24S
del25
del27
A67V
del69
del70
T95I
G142D
del143
del144
del145
N211I
del212
V213G
ins214EPE
G339D
S371F
S371L
S373P
S375F
T376A
D405N
R408S
K417N
N440K
G446S
S477N
T478K
E484A
Q493R
G496S
Q498R
N501Y
Y505H
T547K
H655Y
N679K
P681H
N764K
D796Y
N856K
Q954H
N969K
L981F
BA.1
BA.2
WHU1
WHU1 (RBD only)
WHU1 (delRBD)
S2X35
C002
C144
LyCoV555
REGN10933
REGN10987
C110
C135
S309
S2H97
2-43
N3-1
ACE2-Fc
B
N3-1 HC
N3-1 LC
WHU1 RBD
BA.1 RBD
D105
R106
P373
F375
L371
C
(BA.2) - 1
(BA.1) - 1
Q493R
Q493R
S477N
K417N
S2X35
C002
C144
LyCoV555
REGN10933
REGN10987
C110
C135
S309
S2H97
2-43
N3-1
hACE2
D
E
(BA.1) - 1
(BA.2) - 1
WHU1 + 1
Q498R
N501Y
Q498R
N501Y
F
G
S375F
E484A
E484A
Q493R
E484A
E484A
H
WHU1
WHU1 + Q498R
WHU1 + N501Y
BA.1
BA.1 - Q498R
BA.1 - N501Y
Response (nm)
Time (s)
1500
2000
2500

**Figure S3**: Omicron RBDs broadly escape mAbs while preserving hACE2 affinity

A. Relative binding of RBD-targeting mAbs to WHU1 spikes containing BA.1- and BA.2- mutations. Red, decreased binding; blue, increased binding; relative to the WHU1 spike.
B. Co-structure of N3-1[38] Fab (light chain, pink; heavy chain, violet) in complex with WHU1 RBD (gray) and BA.1 RBD (white), superimposed. Insert shows zoomed in image of large structural rearrangement in 366-377 hairpin loop of BA.1 RBD and the proximity to N3-1's HC residues.
C. Comparison of binding measurements for single amino acid reversions back to the reference (WHU1) amino acids in the BA.1 and BA.2 spikes. Mutations with equal antigenic effects in both spike contexts fall on the diagonal line $y = x$.
D. Comparison of the hACE2 binding caused by adding (+1 amino acid substitution) BA.1-RBD mutations to the WHU1 spike versus reverting (-1 amino acid substitution) the corresponding mutations from the BA.1 spike. Mutations with equal antigenic effects in both spike contexts fall on the diagonal line $y = -x$.
E. Comparison of the hACE2 binding caused by adding (+1) BA.2-RBD mutations to the WHU1 spike versus reverting (-1) the corresponding mutations from the BA.2 spike. Mutations with equal antigenic effects in both spike contexts are expected to fall on the diagonal line $y = -x$.
F. Subset of data from Figure 3E showing mAbs with quaternary binding modes.
G. Subset of data from Figure 3F showing mAbs with quaternary binding modes. All data in Figures S3C-G represent mean ± SD of log-transformed values from at least two biological replicates.
H. Biolayer interferometry (BLI) measurements of the on- and off-rate curves for dimeric hACE2-Fc binding spike variants, corresponding to data in Figure 3J.

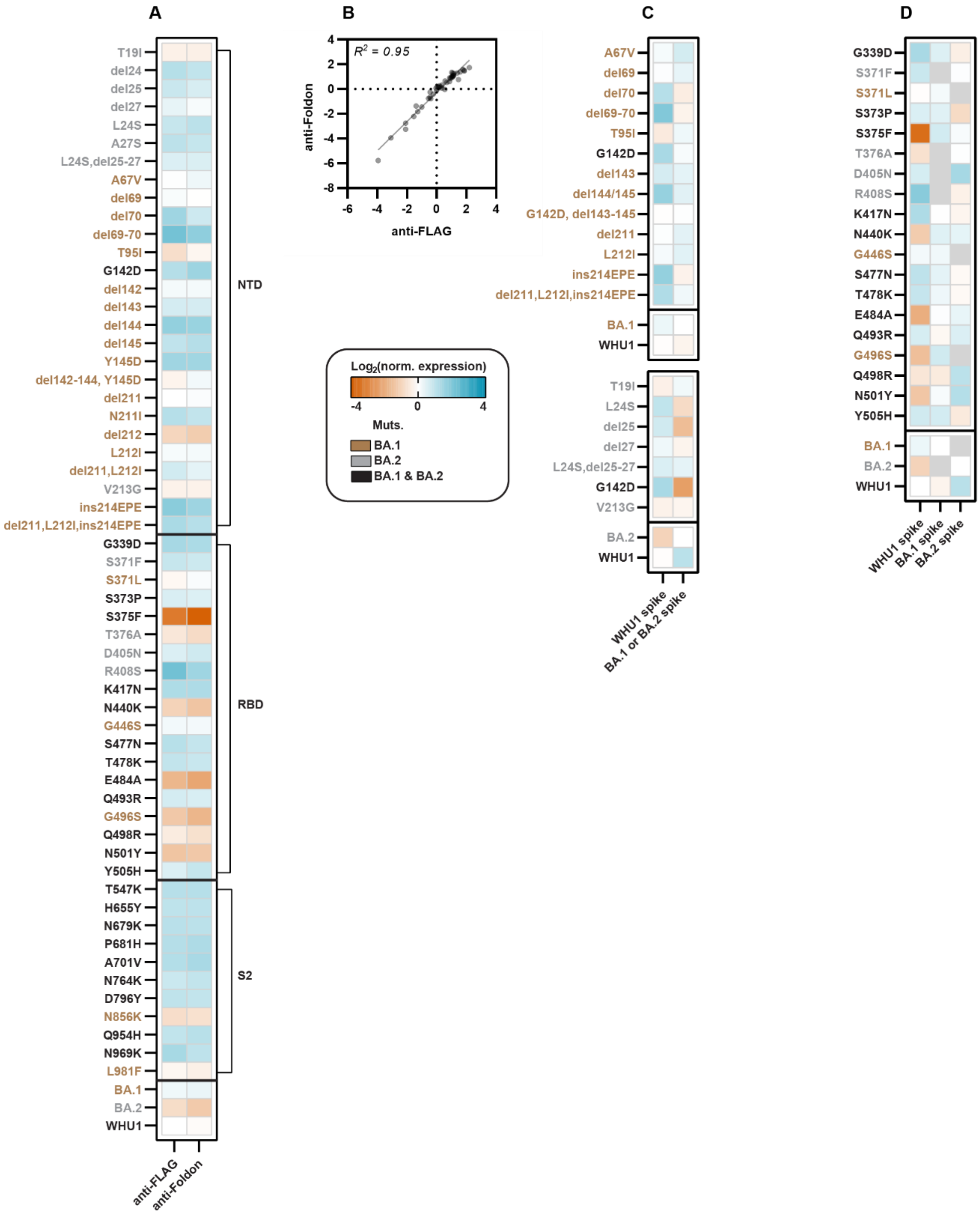
A
B
C
D
T19I
del24
del25
del27
L24S
A27S
L24S,del25-27
A67V
del69
del70
del69-70
T95I
G142D
del142
del143
del144
del145
Y145D
del142-144, Y145D
del211
N211I
del212
L212I
del211,L212I
V213G
ins214EPE
del211,L212I,ins214EPE
G339D
S371F
S371L
S373P
S375F
T376A
D405N
R408S
K417N
N440K
G446S
S477N
T478K
E484A
Q493R
G496S
Q498R
N501Y
Y505H
T547K
H655Y
N679K
P681H
A701V
N764K
D796Y
N856K
Q954H
N969K
L981F
BA.1
BA.2
WHU1
NTD
RBD
S2
anti-FLAG
anti-Foldon
$R^2$ = 0.95
del144/145
G142D, del143-145
Log2(norm. expression)
Muts.
BA.1 & BA.2
WHU1 spike
BA.1 or BA.2 spike
BA.1 spike
BA.2 spike

**Figure S4**: Omicron spike mutations impact spike protein expression

A. Normalized spike protein expression, measured via anti-FLAG or anti-Foldon, for all BA.1 and BA.2 spike mutations in Figures 2B-E, S2H, S2K, and 3B-D. Red, decreased expression; blue, increased expression, relative to the WHU1 spike protein.
B. Comparison of anti-FLAG and anti-Foldon expression measurements for all spike variants in A indicates strong correlation between epitopes. Pearson correlation and linear fit shown.
C. Normalized spike expression values measured for BA.1 (top) or BA.2 (bottom) NTD mutations added to WHU1 spike protein (left column), or reversions of each mutation back to the WHU1 sequence in the BA.1 or BA.2 spikes (right column). Red, decreased expression; blue, increased expression, relative to the WHU1, BA.1 or BA.2 spike proteins, as indicated.
D. Normalized spike expression values measured for BA.1 and BA.2 RBD mutations added to WHU1 spike (left column), or reversions of each mutation back to the WHU1 sequence in the BA.1 spike protein (middle column) or BA.2 spike (right column). Red, decreased expression; blue, increased expression, relative to the WHU1, BA.1 or BA.2 spike proteins, as indicated.

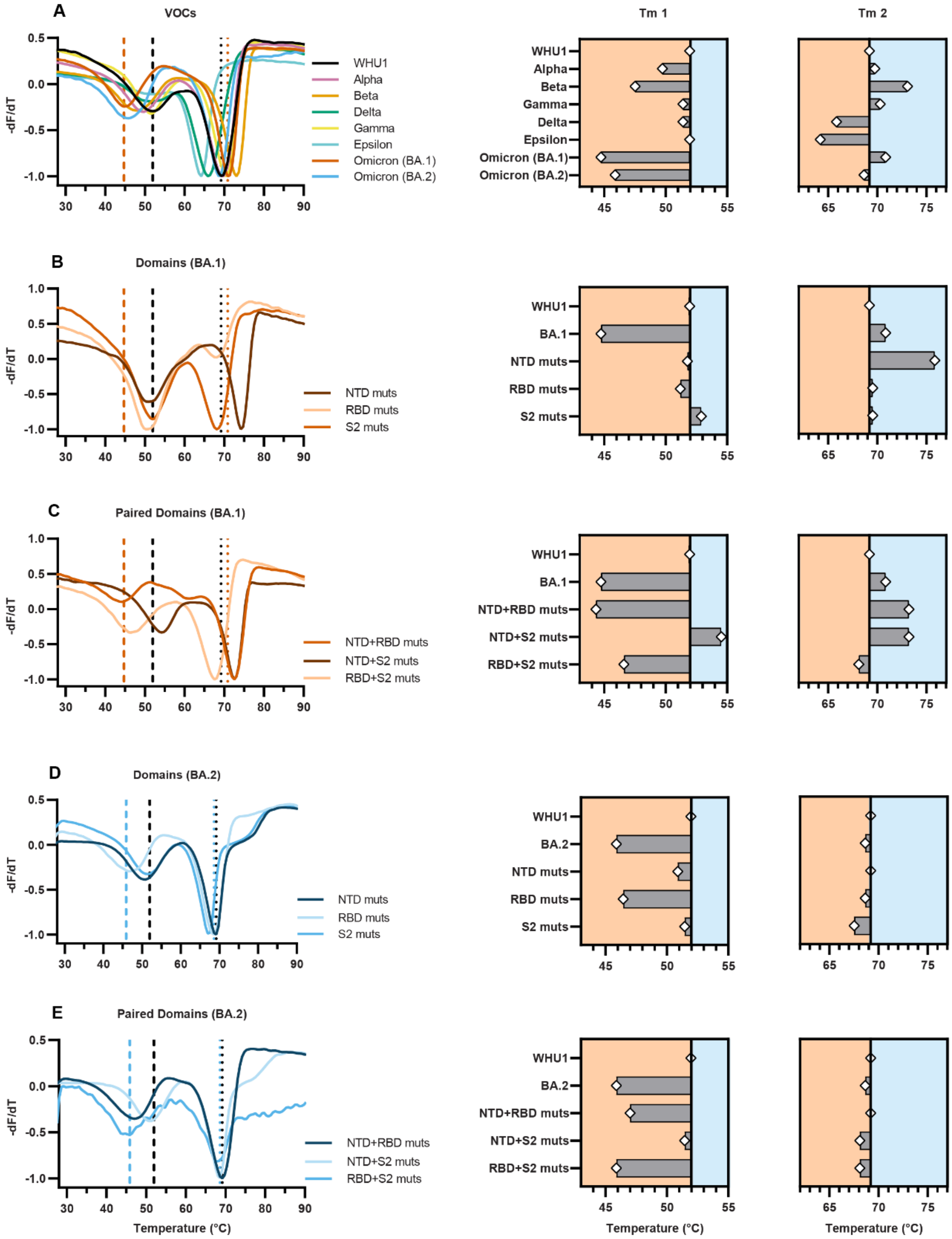

A
VOCs
-dF/dT
WHU1
Alpha
Beta
Delta
Gamma
Epsilon
Omicron (BA.1)
Omicron (BA.2)
Tm 1
Tm 2
B
Domains (BA.1)
NTD muts
RBD muts
S2 muts
BA.1
C
Paired Domains (BA.1)
NTD+RBD muts
NTD+S2 muts
RBD+S2 muts
D
Domains (BA.2)
BA.2
E
Paired Domains (BA.2)
Temperature (°C)

**Figure S5**: Thermostability measurements of spike protein variants show destabilized Omicron RBD

A. Differential scanning fluorimetry (DSF) analysis of spike protein thermostability (see Methods). First and second melting temperatures, $Tm_1$ and $Tm_2$, for WHU1 (black) and BA.1 (red) shown as dotted lines. $Tm_1$ and $Tm_2$ reported for each VOC and VOI (right panels). All values shown relative to WHU1, black vertical line.
B. DSF curves (left) and apparent Tm's (right) for WHU1 spike proteins containing BA.1-NTD, -RBD, or -S2 mutation sets.
C. DSF curves (left) and apparent Tm's (right) for WHU1 spikes containing pairs of BA.1-NTD, -RBD, or -S2 mutation sets.
D. DSF curves (left) and apparent Tm's (right) for WHU1 spike proteins containing BA.2-NTD, -RBD, or -S2 mutation sets. First and second melting temperatures, $Tm_1$ and $Tm_2$, for WHU1 (black) and BA.2 (blue) shown as dotted lines.
E. DSF curves (left) and apparent Tm's (right) for WHU1 spike proteins containing pairs of BA.2-NTD, -RBD, or -S2 mutation sets.
